# 3D *in vitro* modeling of the exocrine pancreatic unit using tomographic volumetric bioprinting

**DOI:** 10.1101/2023.01.23.525142

**Authors:** Viola Sgarminato, Jorge Madrid-Wolff, Antoine Boniface, Gianluca Ciardelli, Chiara Tonda-Turo, Christophe Moser

## Abstract

Pancreatic ductal adenocarcinoma (PDAC) is the most frequent type of pancreatic cancer, one of the leading causes of cancer-related deaths worldwide. The first lesions associated with PDAC occur within the functional units of exocrine pancreas. The crosstalk between PDAC cells and stromal cells plays a key role in tumor progression. Thus, i*n vitro*, fully human models of the pancreatic cancer microenvironment are needed to foster the development of new, more effective therapies. However, it is challenging to make these models anatomically and functionally relevant. Here, we used tomographic volumetric bioprinting, a novel method to fabricate three-dimensional cell-laden constructs, to produce a portion of the complex convoluted exocrine pancreas *in vitro*. Human fibroblast-laden gelatin methacrylate-based pancreatic models were processed to reassemble the tubuloacinar structures of the exocrine pancreas and, then human pancreatic ductal epithelial (HPDE) cells overexpressing the KRAS oncogene (HPDE-KRAS) were seeded in the acinar lumen to reproduce the pathological exocrine pancreatic tissue. The growth and organization of HPDE cells within the structure was evaluated and the formation of a thin epithelium which covered the acini inner surfaces in a physiological way inside the 3D model was successfully demonstrated. Interestingly, immunofluorescence assays revealed a significantly higher expressions of alpha smooth muscle actin (α-SMA) vs. actin in the fibroblasts co-cultured with cancerous than with wild-type HPDE cells. Moreover, α-SMA expression increased with time, and it was found to be higher in fibroblasts that laid closer to HPDE cells than in those laying deeper into the model. Increased levels of interleukin (IL)-6 were also quantified in supernatants from co-cultures of stromal and HPDE-KRAS cells. These findings correlate with inflamed tumor-associated fibroblast behavior, thus being relevant biomarkers to monitor the early progression of the disease and to target drug efficacy.

To our knowledge, this is the first demonstration of a 3D bioprinted portion of pancreas that recapitulates its true 3-dimensional microanatomy, and which shows tumor triggered inflammation.

## 2. Introduction

Pancreatic cancer represents one of the leading causes of cancer-related death worldwide, with a five-year survival rate below 10%^1–3^. Among all types of known pancreatic cancer subtypes, pancreatic ductal adenocarcinoma (PDAC) is the most frequent, accounting for 93% of cancers arising from the pancreas^4^. The absence of clear symptoms in the first stages of PDAC evolution reduces the chances to make an early diagnosis, resulting in a poor clinical prognosis. Indeed, only approximately 10% of the patients are eligible for surgical resection in combination with adjuvant and/or pre-operative therapy, since the majority of cases present spread metastases and extended lesions at diagnosis^5, 6^. Moreover, the unique bioarchitecture of the pancreatic tumor microenvironment (TME) weakens the effectiveness of the current treatments that, despite the advances in the discovery of new therapeutic strategies^7^, result insufficient to treat this particularly aggressive pathology^5, 8, 9^.

In detail, the PDAC microenvironment is composed of approximately 90% desmoplastic stroma consisting of collagen, fibronectin, fibrillar collagen and hyaluronic acid (HA)^10^. This dense stromal tissue creates a hypoxic environment and plays a key role in disease progression and drug resistance, constituting a barrier which impedes therapeutics access. The stroma arises from the excessive extracellular matrix (ECM) deposition by pancreatic stellate cells (PSCs) that are responsible for the intense desmoplastic reaction occurring within the tissue surrounding the cancer cells^11, 12^. More precisely, in healthy tissue PSCs surround the epithelial cells that constitute the pancreatic functional unit (Fig. 1a_i_), responsible for the secretion of the digestive enzymes. It is in this region that pancreatic intraepithelial neoplasia (PanIN) develops^5, 13, 14^. PanIN is the early precursor lesion which progresses to the development of PDAC through mutation of specific genes in epithelial cells (Fig. 1a_ii_).

**Fig. 1.**
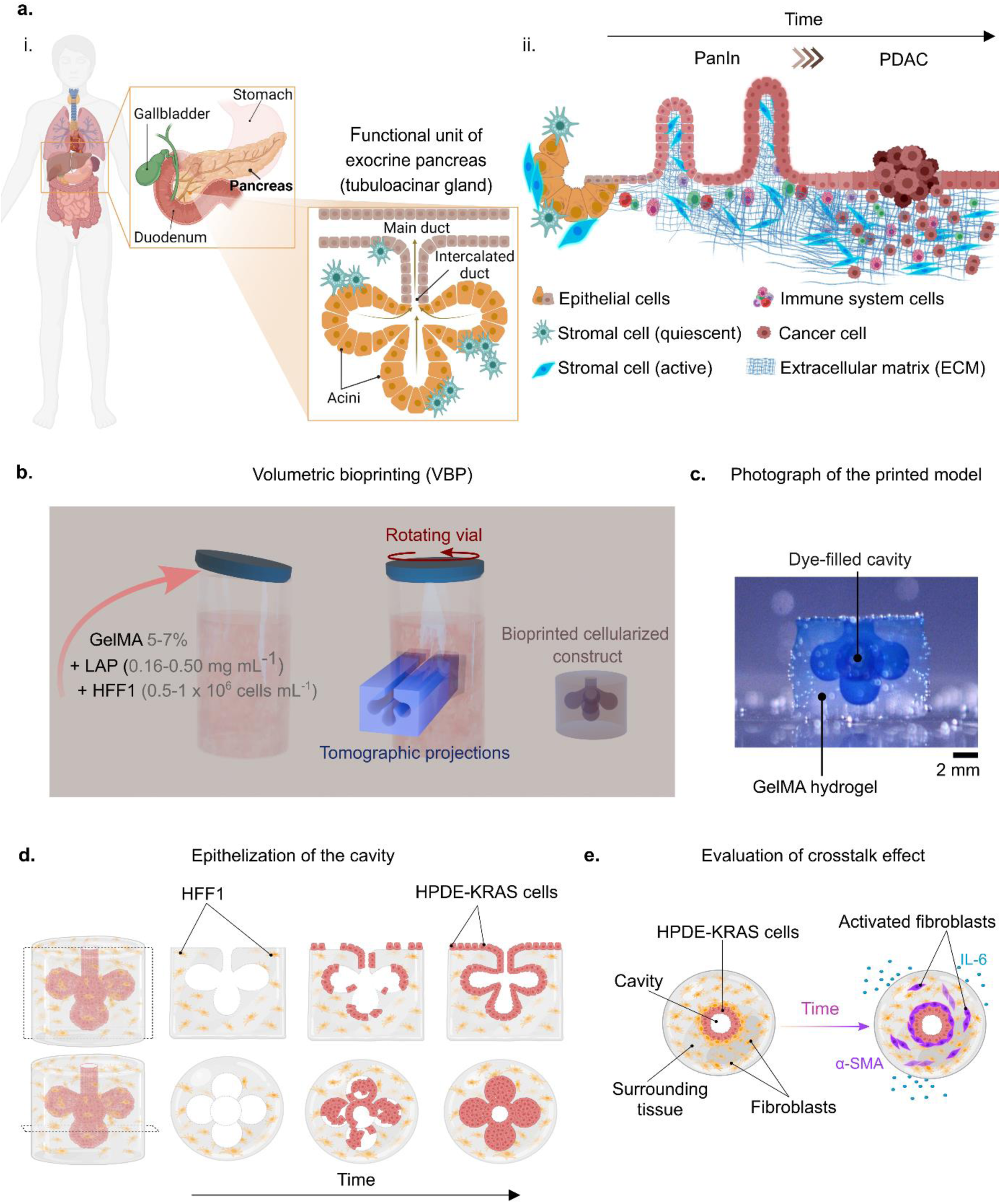
Experimental pipeline for tomographic volumetric bioprinting of pancreatic cancer models. **a**. Pancreatic cancer typically occurs within the functional unit of exocrine pancreas, composed of epithelial cells surrounded by pancreatic stellate cells (PSCs) (i). Schematic representation of pancreatic intraepithelial neoplasia (PanIN) evolution into mature pancreatic ductal adenocarcinoma (PDAC) inspired from Liot et al.^15^. Particular focus is made on the crosstalk between PSCs and PDAC cells which promotes tumor evolution (ii). **b.** In tomographic volumetric bioprinting, a gelatin methacryloyl (GelMA) solution in cell medium containing the photoinitiator lithium phenyl-2,4,6-trimethylbenzoylphosphinate (LAP) and human fibroblasts (HFF1) is poured into glass vials and printed. A cell-laden construct that mimics the functional unit of exocrine pancreas is obtained. **c**. Photograph of a bioprinted construct (immersed in water) with the cavity filled with a glycerol-based blue dye (for visualization purpose only). **d**. Human pancreatic ductal epithelial (HPDE) cells overexpressing the KRAS oncogene (HPDE-KRAS cells), are injected into fibroblast-laden bioprinted constructs, where they attach and coat the acini inner surfaces with time. **e.** To recapitulate fibroblast-associated inflammation or activation, the interaction between HPDE-KRAS cells and the surrounding fibroblasts is monitored by measuring the expression of α-SMA versus actin in the cytoskeletons of the latter and the release of IL-6 cytokines in the supernatants. (Figure drawn using Biorender.com).

Although the alterations that give rise to PanIN precursor lesions are still to be clarified, hallmarks associated to pathology onset have been identified. Indeed, the oncogene KRAS in the epithelial cells have been reported as the more frequent mutation leading to the progression of PanIN^15^. Besides mutation in epithelial tissue genome, PDAC formation is also characterized by the desmoplastic reaction induced by cancer cells. Therefore, during the pancreatic carcinogenesis, the PSCs, that are in a quiescent state and exhibit a star-shaped morphology, activate in response to inflammatory cues and cancer cells-derived factors, acquiring a myofibroblasts-like phenotype which is characterized by the increased assembly of alpha-smooth muscle actin (α-SMA) stress fibers^16, 17^ (Fig.1b). Typically, activated PSCs assemble in a core-shell like structure surrounding the cancer cells and start to interact with them by generating a complex autocrine and paracrine signaling interplay^18, 19^. In this intricate framework, the stromal components actively interact with pancreatic cancer cells through different ways that significantly affect gene expression patterns, metabolic activities, invasion/metastasis phenomena and resistance mechanisms^20^. In particular, the activated PSCs released cytokines such as interleukin 6 (IL-6) and growth factors (*e.g.*, TGF-β), leading to an increase in inflammatory cues fostering the mutation of the oncogene KRAS in the epithelial cells and the progression from PanIN to PDAC^18, 21^ (Fig. 1a_ii_).

The understanding of dynamic phenomena involved in PDAC-stroma crosstalk might expand the knowledge of the pathology and consequently discover innovative biological targets. Indeed, even though important risk factors can contribute to the development of the pancreatic cancer (like smoking, obesity, type 2 diabetes, chronic pancreatitis, and alcoholism^22^) and although the mechanisms of evolution from the neoplasia are well documented^23, 24^, the alterations causing the early lesions still remain unclear^25^. Therefore, the understanding of pancreatic cancer raises interests among the scientific community, currently developing efficient PDAC *in vitro* models in order to detect the disease earlier and design effective therapies thus improving patients’ prognosis^26–31^. Although recent works have shown the possibility of modeling the PDAC microenvironment *in vitro*^26, 27, 29, 32–37^, the tumor-stroma interplay remains arduous to replicate and monitor in functionally effective models^38–41^. Heterotypic 3D spheroids, patient-derived organoids, cancer-on-a-chip platforms, and 3D biofabricated constructs are the currently available bioengineered 3D models mimicking the pancreatic tumor-stroma interplay^19^. However, only a few novel studies in literature focus on the development of biomimetic platforms reproducing the microanatomy (in terms of 3D architecture and cellular composition) of the exocrine pancreas and lack to resemble the native compartmentalized architecture of tumor microenvironment that is widely recognized to affect cell functionality and cancer-cell response to therapeutics^42–44^. In particular, the gland complex geometry has been reproduced in simplified ways by employing different techniques such as viscous fingering and extrusion-based methods^45–48^, which result in low reproducibility, throughput and shape fidelity. These limitations can be overcome using volumetric bioprinting (VBP), an emerging light-based technology capable of fabricating 3D constructs with high-resolution and complex geometries rapidly^49–52^. Indeed, this technique permits to print hollow structures without the need for support structures and in a very short building time (down to a few tens of seconds compared to tens of minutes for layer-by-layer approaches)^50^. Furthermore, one of the main advantages of VBP is the cell-friendly procedure relying on the one-step manufacturing process which reduces the stress experienced by cells as compared to other multistep techniques such as the common solvent-casting method^53^. More specifically, VBP consists of illuminating a photosensitive cell-laden hydrogel with visible light from multiple angles, using a sequence of tomographic back projections of the desired object^54–56^, leading to the photopolymerization of the material. We adopt VBP to develop a fully human 3D *in vitro* model resembling the physiological tubuloacinar gland morphology of the exocrine pancreatic unit^57, 58^ (Fig. 1b). In particular, a gelatin methacrylate hydrogel (GelMA) has been *ad hoc* prepared and loaded with human fibroblasts (stromal cells) to mimic the stromal compartment. Fibroblast-laden structures were fabricated by VBP to mimic the tubuloacinar physiological architecture (Fig. 1b,c). Then, we introduced human pancreatic ductal epithelial cells stably expressing the KRAS oncogene (HPDE-KRAS) inside the construct’s cavity and monitored the co-culture overtime (Fig. 1e). We analyzed the tumor-stroma crosstalk effect measuring the appearance of a myo-fibroblast phenotype, by quantifying expression of α-SMA proteins in fibroblasts and evaluating the release of inflammatory cytokines (Fig. 1f).

## 3. Results

### 3.1 Fabrication of exocrine pancreatic units through VBP

The constructs initially fabricated as proof of concept followed a ductal geometry, with an acinus of larger diameter at the end (sup. video 1). This geometry is challenging to fabricate with conventional methods and without employing support structures or joining two compartments. In VBP, we projected a set of tomographic light patterns into cell-laden GelMA (5% w/v in DMEM w/o phenol red + 0.5 million fibroblasts mL^-1^) in which we added the photoinitiator at a low concentration (0.16 mg mL^−1^) (Fig.2). The selected concentration of the photoinitiator allows the crosslinking of GelMA upon visible light irradiation at 405 nm wavelength (Fig. 2a,b) and it is low enough not to induce cytotoxicity and light absorption. The photorheological test proved the sol-gel transition of GelMA as a result of light irradiation (Fig. 2b). The increment of storage modulus (G’) until reaching a stable plateau at 0.4 kPa indicates the elastic response of the material, whose viscoelastic properties are comparable to that of pancreatic tissue^59^ (sup. fig. 1). Moreover, the GelMA hydrogels have a swelling capacity of 726 ± 27 % (sup. table 1), suggesting a high porosity of the hydrogel network resulting in a good level of medium diffusion within the gel^60, 61^. The light-scattering effect caused by the presence of cells within the gel causes a loss of resolution in the constructed object, such as a resulting obstructed duct or an incomplete acinus. However, by performing a numerical correction of the light dose that uses quantitative information on the light scattering^50^, we obtained an improvement in shape fidelity and resolution of the printed structures (Fig. 2c,d). This correction works by measuring the light scattering profile of the cell-laden hydrogel and then updating the projected tomographic patterns accordingly so that light reaches the desired volume and not elsewhere in the vial. In a second stage, we furtherly optimized the VBP process to improve the model complexity, by reducing the size and introducing more acini (five) within the same hydrogel construct. We obtained small complex structures which reproduce the tubuloacinar gland morphology of the physiological exocrine pancreatic unit with the outer cylinder and the inner acinus diameters rescaled relative to the proof-of-complex structure by a factor of 1.4 and 3, respectively (Fig.2e, sup. video 2). To fabricate these constructs 7% w/v GelMA in DMEM w/o phenol red containing 1 million fibroblasts mL^−1^ and 0.5 mg mL^−1^ LAP was used. Features of the final geometry, such as the wall thickness and the duct diameter, were optimized to guarantee printability and maximize anatomical relevance. This is because there is a trade-off between structural integrity and cell viability, as metabolic activity in cells within hydrogels is known to decrease with distance from the outer borders of the hydrogel^62^. Fig. 2f shows the effectiveness of VBP in reproducing the stromal component surrounding the acinar structures in the exocrine tissue. Indeed, fibroblasts formed a natural network within the hydrogel and around the acini (Fig. 2f_i_) while they are not present in the hollow parts of the construct (Fig. 2f_ii_).

**Fig. 2.**
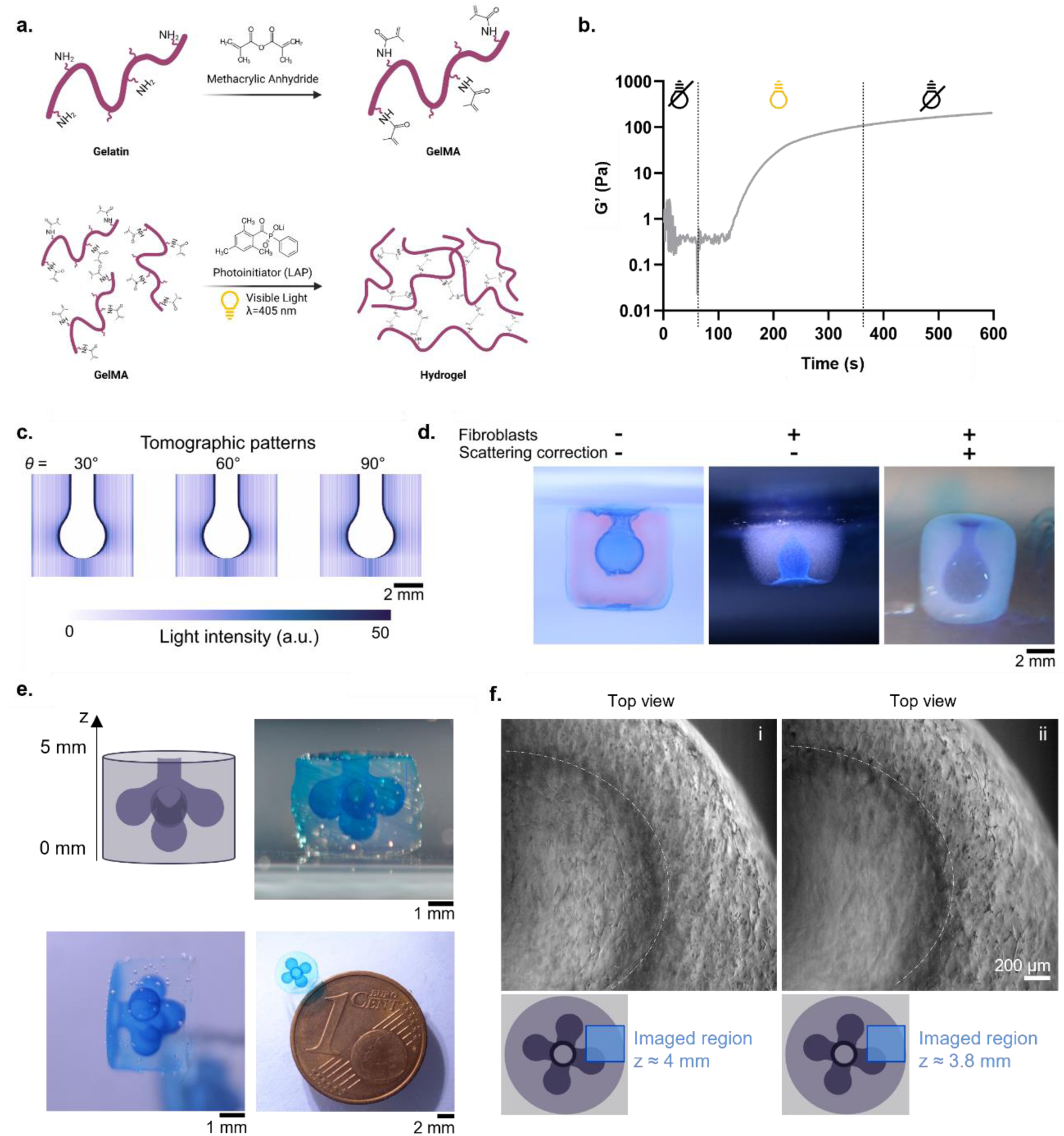
Fabrication of viable 3D pancreatic ductal models. **a.** Synthesis and photocrosslinking of gelatin methacryloyl (GelMA) inspired from Yoon et al.^63^. Gelatin was reacted with methacrylic anhydride (MA) to introduce a methacryloyl substitution group on the reactive amine and hydroxyl groups of the amino acid residues. The GelMA photocrosslinking occurs after exposure to visible light (405 nm wavelength). The free radicals generated by the photoinitiator initiate the chain polymerization with methacryloyl substitution resulting in the hydrogel formation. **b.** Photorheology test to evaluate the photocrosslinking kinetics by monitoring the storage modulus (G’) overtime. The crosslinking reaction started after 60 s following the emission of visible light (405 nm wavelength). **c.** Some tomographic patterns used to fabricate the scattering-corrected constructs. In total, 1000 different tomographic patterns are displayed along each turn of the cylindrical vial during printing. **d.** Photographs of 3D printed GelMA constructs, immersed in water and with the duct filled with a glycerol-based blue dye for illustrative purposes. Including cells in the gel affects print fidelity, but it can be compensated by correcting for scattering effects. **e.** Photographs of the new model, with enhanced level of geometrical complexity and biomimicry compared to the proof-of-concept structure constituted by a single acino-ductal cavity. **f.** Optical microscopy brightfield images of the fibroblast-laden multiacinar construct obtained by VBP. The images show a portion of one of the 5 acini at two different focal planes. Schematics show the region where these microscopy images were acquired.

### 3.2 Viability of HFF1 inside the bioprinted exocrine pancreatic unit

We monitored the state of fibroblasts within the bioprinted exocrine pancreatic unit models first, to assess their viability and to identify the optimal temporal window for establishment of the co-culture. Cell proliferation occurred over time after 72h as confirmed by the metabolic activity of fibroblasts which increased with time. Indeed, Fig. 3a shows fluorometric measurements of the CellTiter-Blue cell-viability assay (*n_GelMA + cells_* = 7, *n_GelMa_* = 7), in which resazurin is reduced by metabolic reactions in the cells to resorufin, a fluorescent molecule. Higher fluorescence intensity indicates higher cell viability. A significant (p < 0.0001), marked increase in metabolic activity from 72 to 96 hours after printing was observed, which is sustained at least until 9 days after printing. This significant increment in cell viability at 96h can be ascribed to the cell recovery from stress that the biofabrication process may cause suggesting an optimal time window for seeding of epithelial cells starting from 4 days after the bioprinting process. Furthermore, the fluorescence intensity using bioprinted GelMA hydrogels without cells was also measured as GelMA hydrogels have residual free radicals left after gelation, which can themselves account for the reduction of rezasurin^64^. No significant fluorescent intensities were measured in control condition as values were 2 orders of magnitude lower than the one in cell-laden hydrogels (Fig. 3a).

**Fig. 3.**
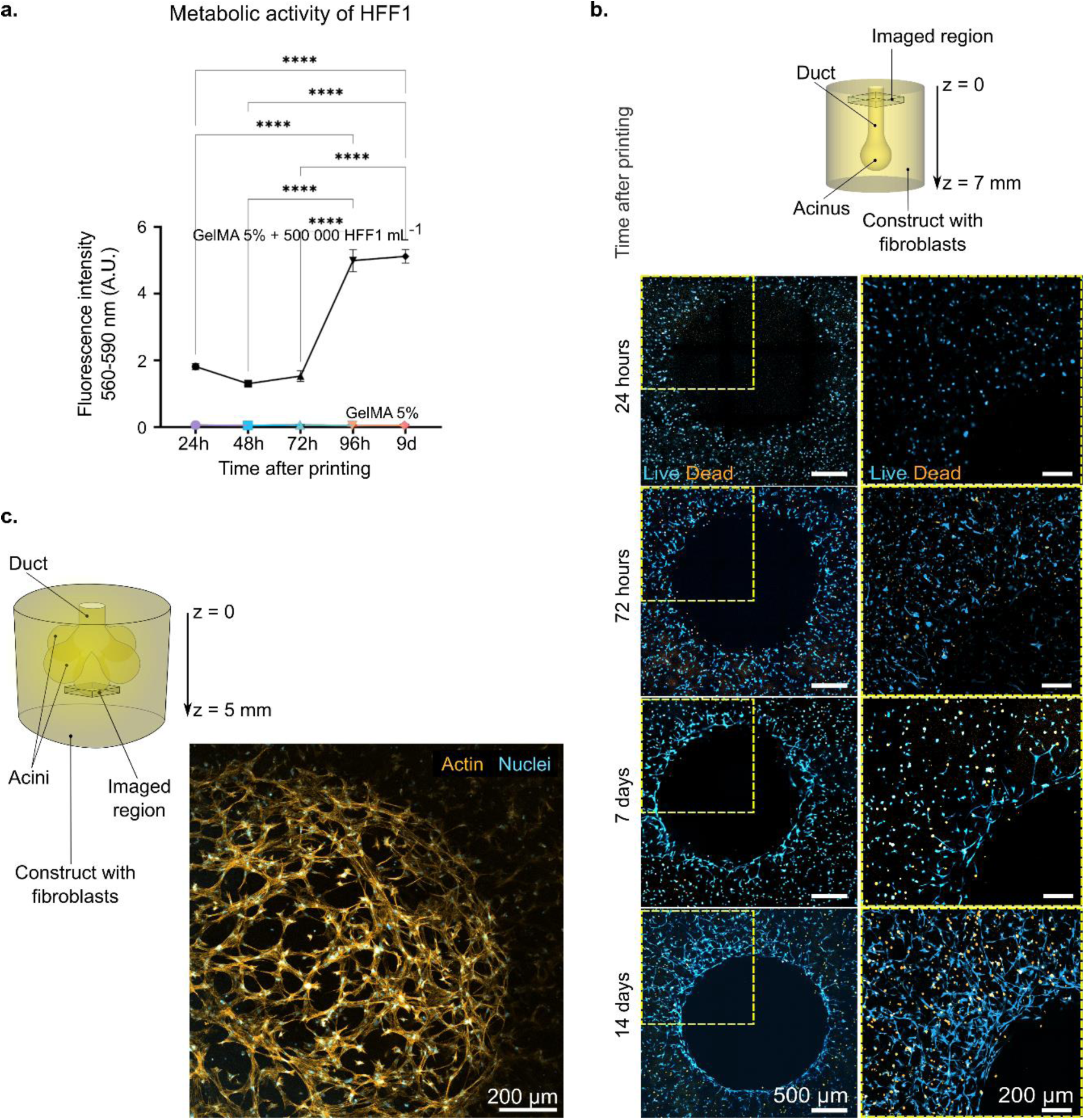
Viability of pancreatic ductal models. **a.** Metabolic activity of fibroblasts (black) as a function of time, measured from the reduction of resazurin. Cell-free printed hydrogels (color) are used as a control (n_GelMA + cells_ = 7, n_GelMA_ = 7, three measurements per sample). Tukey’s multiple comparisons test: \**p < 0.05*, \*\**p < 0.01*, \*\*\**p < 0.001*, \*\*\*\**p < 0.0001*. Error bars represent one standard deviation. **b.** Live/Dead assay of fibroblasts performed on different samples 1, 3, 7 and 14 days after volumetric bioprinting. A schematic shows the region where these microscopy images were acquired. **c.** Representative high-magnification multi-channel fluorescence microscopy image of the bottom acinus surface 12 days after the volumetric bioprinting process. A diagram illustrates the area from which the z-stack microscopy image was acquired.

The distribution of viable cells within the GelMA constructs was also monitored by fluorescence microscopy.

Fig. 3b shows fluorescence microscopy images of constructs 24 hours, 72 hours, 7 days, and 14 days after printing. Live cells, shown in blue, had been stained with calcein-AM, a membrane-permeant dye that is converted into a fluorescent calcein by intracellular esterases. Dead cells, shown in orange, had been stained with ethidium homodimer-1, which is a membrane-impermeant high-affinity nucleic acid stain that is weakly fluorescent until bound to DNA. The micrographs show the region around the duct, 300 μm deep inside the constructs, with fibroblasts assuming an elongated shape and fully colonizing the inner walls of the duct over time. Images also demonstrate that most cells survived after the printing process and that they were homogenously distributed within the hydrogels at 24h. Although there are some dead cells in the hydrogels starting from 1 week after the bioprinting process, live fibroblasts grow with time, exhibiting an elongated shape, characteristic of healthy cell states (sup. fig. 2). In Fig. 3c a representative z-stack multi-channel fluorescence microscopy image, which correspond to the intensity sum over planes along 480 μm of the microscope’s optical axis, of the bottom acinus of a fibroblast-laden construct (12 days after printing process) shows HFF1 assuming their characteristic elongated shape within the bioprinted structure. In particular, this acquisition proves that stromal cells formed a highly biomimetic concave network which surrounds the 3D acinar cavity (sup. video 3).

### 3.3. Colonization of the acini inside the VBP construct by HPDE-KRAS cells

The possibility to achieve a compartmentalized structure recreating the cell distribution found *in vivo* was assessed by seeding HPDE-KRAS cells within the cavity of the bioprinted structure. To this end, the capacity of human pancreatic ductal epithelial cells to epithelize the walls of the cavity (duct and acini) of the bioprinted constructs was evaluated by means of immunofluorescence imaging. At first, we injected HPDE-KRAS cells at different densities into the GelMA bioprinted structures without fibroblasts. Results indicated that cells seeded at a lower density (1400 cells mm^-2^; cell number ratio 0.9:1 HPDE-KRAS:HFF1) formed cell clusters after 3 days and subsequently epithelialized most of the cavity within 3 weeks. In contrast, the same tendency was not observed with HPDE-KRAS cells seeded at a higher density (7200 cells mm^-2^) (sup. fig. 3). Indeed, a higher number of cells resulted in cell sedimentation, distinguishable by dark spots in brightfield microscope images, at the bottom of the acini within 3 days of seeding (sup. fig. 3_ii, iv_). However, epithelial cells were observed to migrate from the bottom of the acini and colonize the lateral and more vertical parts of the cavity, ultimately forming a thin epithelium after 7 days (around 2 weeks faster than cells seeded at a lower density). Therefore, we used the same cell density (7200 cells mm^-2^; cell number ratio 4.6:1 HPDE-KRAS:HFF1) to seed HPDE-KRAS cells within the cavity of HFF1 bioprinted structures 5 days after the bioprinting process. This time point was selected for implementing the co-culture conditions to ensure HFF1 recovery during the first 96h, when the viability significantly increased (Fig. 3b).

Seeded HPDE-KRAS cells colonized the multiple acini of acellular and fibroblast-laden hydrogels (Fig. 4a). These epithelial cells remained viable up to 21 days after seeding (Fig. 4b) and packed themselves densely into epithelia of up to 11 300 cells mm^-2^ (Fig. 4c). After culturing them for several days, we verified that they colonized the full extent of the convoluted geometry of the bioprinted constructs (Fig. 4d). High coverage was achieved faster by HPDE-KRAS cells growing on fibroblast laden hydrogels, as seen in Fig. 4e and sup. video 4, probably thanks to the secretion of extracellular matrix components by fibroblasts. The resulting constructs included densely epithelized acini surrounded by fibroblasts (Fig. 4e,g). We verified the expression of E-cadherin, a cell–cell adhesion glycoprotein, by these epithelial cells. E-cadherin is a hallmark of intercellular adhesion, particularly in growing epithelia^65^. Light sheet and confocal fluorescence microscopy images revealed that HPDE cells in the bioprinted constructs expressed E-cadherin at higher rates than fibroblasts, as expected, and particularly in proliferating and colonizing cells, compared to the cells attached to the cavity‘s surface in a confluent state (Fig. 4d,e,f). Remarkably, we observed a marked polarity in the expression of actin and E-cadherin in established epithelial cells (14 days after seeding), as seen in Fig 4h. E-cadherin expression was higher towards the lumen of the cavities, away from the hydrogel; while actin expression was higher towards the outside, in the section of the epithelium in contact with the hydrogel. The resulting epithelia were thin, at 37 ± 1.5 µm, and were composed of 1 to 3 layers of cells in general (sup. fig. 4 shows the separate fluorescence channels, including DAPI, where nuclei are visible).

**Fig. 4.**
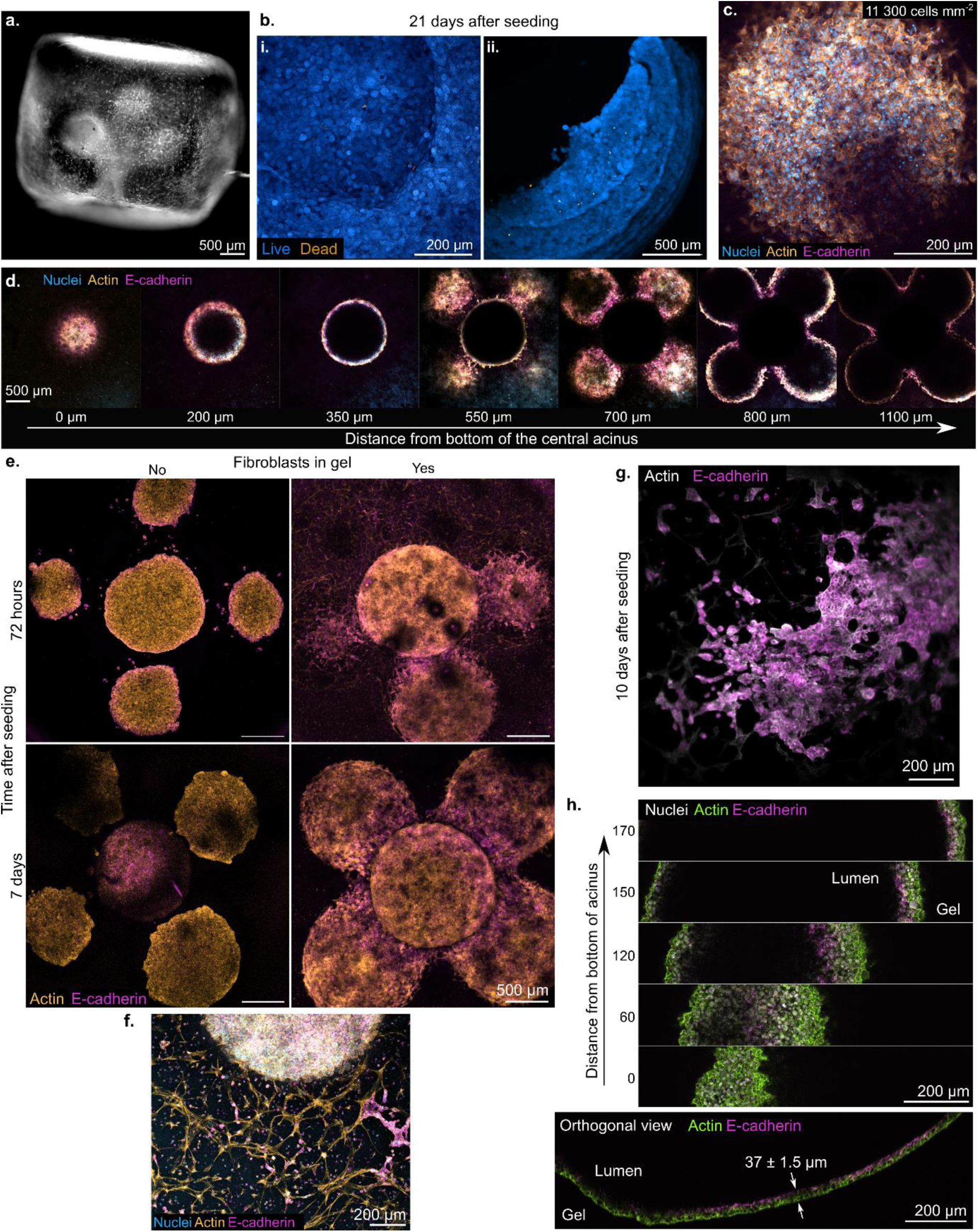
Epithelization of multi-acinar constructs. **a.** Dark-field photograph of a hydrogel containing fibroblasts and HPDE-KRAS cells in the acini (visible as a white cloud). **b**. Live/Dead staining of the ductal epithelial cells 21 days after having been seeded in the construct. Top view (i) and oblique view (ii) of one of the spherical acini. **c**. The seeded ductal epithelial cells form dense linings of up to 11 300 cells mm^-2^ (72 hours after seeding), as seen in this maximum intensity projection of multi-channel confocal images. **d.** Slices from confocal microscopy images of an entire construct containing fibroblasts in the gel and ductal epithelial cells (+ KRAS oncogene), 7 days after seeding them. **e.** Coverage of the epithelium increases with time and is more complete when HPDE-KRAS cells are co-cultured with fibroblasts, as seen from these maximum intensity projections of confocal microscopy images. **f.** Epithelial cells express E-cadherin, a cell–cell adhesion glycoprotein, when growing into epithelia. Maximum intensity projections of light sheet microscopy images showing a curved vertical section of the construct. Fibroblasts are indicated with green arrows. Gravity pulls towards the bottom of the image. **g.** Human pancreatic ductal epithelial cells form a dense epithelium (circular white structure in the upper part of the image) 7 days after having been seeded into a gel containing fibroblasts (1 x 10^6^ cells mL^-1^). Seeding was done 5 days after printing. **h**. Spatial distribution of E-cadherin vs. actin expression in ductal epithelial cells, from high-resolution confocal fluorescence microscopy images. The orthogonal view of this set of images was used to measure the average thickness of the resulting epithelium at 37 ± 1.5 µm. (n = 5 measurements, with standard error of the mean). Gravity pulls towards the bottom of the orthogonal view. From HPDE-KRAS in gels without fibroblasts, 14 days after seeding.

### 3.5 Evaluation of cell-cell crosstalk within the VBP model

The effect of crosstalk between HPDE cells and HFF1 was monitored by evaluating changes in the cytoskeleton composition of the latter. In particular, the activation of fibroblasts was monitored in co-culture with healthy HPDE (HPDE-WT) and HPDE overexpressing the KRAS oncogene (HPDE-KRAS) using a single acinus design to allow more precise monitoring and imaging. Healthy HPDE (HPDE-WT) and HPDE overexpressing the KRAS oncogene (HPDE-KRAS) were seeded in the fibroblast-laden hydrogels 4 days after the bioprinting process. Immunofluorescence microscopy was performed on 300 μm thick transversal slices of HFF1-laden constructs containing HPDE-WT cells, HPDE-KRAS, and without HPDE cells as a control (n = 4 replicas for HPDE-WT 72 hours; n = 3 for all other treatments) at 5 and 7 days after printing (24 and 72 hours after HPDE seeding), as seen in Fig. 4a. We used thin slices of the gel obtained with a vibratome to guarantee that antibodies would penetrate evenly throughout the constructs and that all cells in them could be imaged. Anti-α-SMA antibody, phalloidin-FITC (an actin marker), and DAPI (a DNA marker) were used to identify cells morphology and to evaluate the appearance of a myofibroblast phenotype associated with an increased expression of α-SMA (sup. fig. 5). Qualitatively, we saw that fibroblasts in constructs seeded with HPDE-KRAS cells exhibited stronger expression of α-SMA, and that this expression increased with time. We developed an algorithm to automatically and blindly quantify the expression of α-SMA with respect to actin over hundreds of cells in several complete slices of the constructs, which were imaged under a confocal microscope following a standardized protocol. The automated analysis, which utilized the DAPI signal for cell detection, computed the ratio of α-SMA to actin intensities as a proxy for fibroblast activation, as depicted in Fig. 4b. This analysis showed that the mean fibroblast activation increased after exposure to HPDE-KRAS cells, but not so after exposure to the non-cancerous HPDE-WT. The level of fibroblast activation resulted also significantly higher after being in co-culture with HPDE-KRAS for 72 hours compared to 24 hours (p = 0.018). The inflammatory response involved in the crosstalk between stromal and epithelial cancer cells was also analyzed in terms of pro-inflammatory cues produced by fibroblasts (fig. 4c). Results indicated a higher IL-6 release by fibroblasts in co-culture with HPDE-KRAS cells for 48 and 72 hours, in comparison with HFF1 alone or HFF1 under co-culture with healthy HPDE cells (HPDE-WT). The IL-6 level significantly increased after 24h for fibroblasts co-cultured with HPDE-KRAS cells while it remained constant for fibroblasts cultured with healthy epithelial cells. We also studied the dependence of fibroblast activation with the distance to the duct at the single cell level. Fig. 4d shows these measurements, suggesting an increased activation for fibroblasts closer to the duct, where HPDE-KRAS cells laid. In contrast, fibroblasts co-cultured with HPDE-WT cells or with no HPDE cells did not exhibit a decaying degree of activation with distance to the duct. These results indicated that the activation of fibroblasts occurs predominantly when they are co-cultured with HPDE-KRAS cells and that the dependence of α-SMA expression on the distance from the duct is evident only in this condition.

## 4. Discussion

Despite several attempts to understand pancreatic cancer progression over the past decades, pancreatic ductal adenocarcinoma (PDAC) remains one of the most lethal tumors, with the highest 1-year, 5-year and 10-year mortalities of any cancer type^3^. Modeling the dynamic phenomena involved in tumor-stroma interplay is essential not only to better understand the disease but also to develop new therapeutics. Indeed, the stromal tissue surrounding the PDAC site represents a histopathological hallmark of pancreatic cancer^66–71^ and plays a fundamental role in tumor progression^72, 73^.

Therefore, in this study we developed a 3D *in vitro* model of the exocrine pancreas which mimics the compartmentalized architecture of the native tissue and recapitulates the stromal and pancreatic cancer cell crosstalk in the same miniaturized construct. In particular, this work focuses on modeling the tubuloacinar gland to provide the biomimetic architectural stimuli that cells experience *in vivo*. Indeed, as many studies have evidenced, the geometry of the substrate significantly affects cellular behavior^74^. Specifically, surface curvature at multi-cellular level is a key geometrical factor that modulates tissue growth and cell organization, such as cell polarization and orientation^75, 76^. We co-cultured human fibroblasts to model the stromal component and human pancreatic epithelial cells expressing the KRAS oncogene, to reproduce the pathological exocrine pancreatic tissue, also in terms of its morphology (Fig. 1). To achieve these features, the biomimetic gland was fabricated exploiting tomographic volumetric bioprinting. This one-step, cell-friendly and scalable approach allowed us to fabricate 3D cell-laden hydrogels incorporating a duct converging to five acini with high shape fidelity in less than one minute (Fig. 2). We demonstrated the printing of relevant object shapes for biological studies while maintaining a suitable environment for the growth of stromal cells (HFF1) that remained viable and active for at least 2 weeks after the manufacturing process (Fig. 3). This is in line with other works, which have cultured viable tomographically printed constructs for several weeks^49, 54, 77^. Moreover, the results proved the beneficial effects given by GelMA as bioink, matching with previous reports on the extensive use of this material in biomedical applications^78–82^. This innovative biofabrication approach avoids the technical difficulties and time-consuming procedures associated with the assembling of different cellularized compartments into a unique 3D structure^45^. The fabrication of small (< 200 µm) cavities within cellularized hydrogels remains an open challenge in tissue engineering. Future work in volumetric bioprinting should push towards bridging this gap.

The cavity within the printed construct constitutes a biomimetic cavity surrounded by human stromal cells (Fig. 3c) which can be easily epithelialized by seeding the human pancreatic ductal epithelial cells, suspended in a proper volume of cell medium. We monitored the proliferation of HPDE-KRAS cells over time and assessed their ability to cover the inner walls of the lumen as they grew as an epithelium (Fig. 4), as reported by other studies in literature^46, 83^. Additionally, the E-cadherin expression observed in epithelial cells covering the multiacinar cavity closely resembled the epithelial cell polarity observed *in vivo*^65, 84^.

Under co-culture conditions we monitored the activation of stromal cells by quantifying, through a custom-made Python code, the signal intensity coming from the expression of α-SMA proteins (Fig. 5a,b,d). The results, showing a higher α-SMA expression in fibroblasts co-cultured with HPDE-KRAS rather than in contact with HPDE-WT, allowed to validate this *in vitro* model as it can efficiently mirror the physiological inflammation cascade occurring in activated stromal cells^32, 36, 37, 85, 86^. Moreover, the cell crosstalk between stromal and HPDE-KRAS cells within the VBP model was investigated also focusing on fibroblast inflammation mediated by IL-6 cytokines (Fig 5c). The release of IL-6 by inflamed tumor-associated fibroblasts is known to significantly influence the interplay between PDAC and its stroma, regulating various mechanisms such as angiogenesis, epithelial-to-mesenchymal transition, and immunosuppression^87, 88^. Our findings revealed increased IL-6 release by fibroblasts co-cultured with HPDE-KRAS cells for 48 and 72 hours, compared to fibroblasts alone or in co-culture with healthy HPDE cells (HPDE-WT). This observation aligns with existing literature highlighting the role of the KRAS oncogene in driving IL-6 production by stromal cells^89, 90^. Thus, our developed VBP model effectively recapitulates the pathological scenario *in vivo*, wherein fibroblasts secrete IL-6 when inflamed by cancer cells^91^. Interestingly, the higher IL-6 secretion and α-SMA expression in fibroblasts co-cultured with HPDE-KRAS cells in close proximity (Fig. 5d) suggest a potential correlation with a specific cancer-associated fibroblasts (CAFs) subpopulation (csCAFs) which has been lately identified^92^ in the stroma of early-stage PDAC adjacent to tumor cells^93, 94^. However, RNA sequencing analyses should be performed to further investigate this bioactivity in HFF1.

**Fig. 5.**
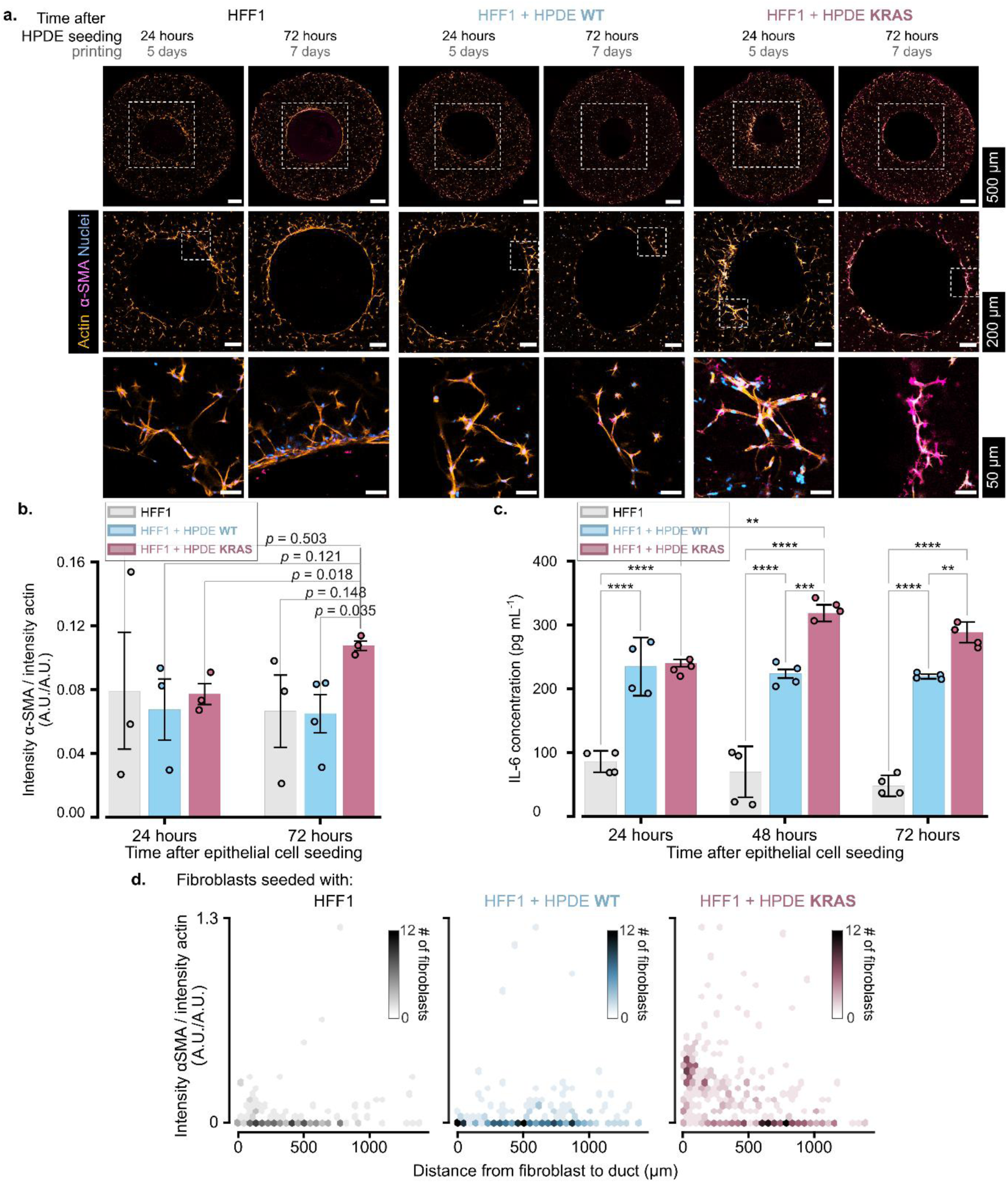
Tomographic biofabricated 3D pancreatic models recapitulate inflammation of cancer-associated fibroblasts. **a.** Fluorescence microscopy images of full slices of constructs without HPDE cells and seeded with HPDE-WT or HPDE-KRAS cells 24 hours (5 days) and 72 hours (7 days) after bioprinting. Lower rows correspond to close-ups of the dashed regions. Scale bars top row: 500 μm, mid row: 200 μm, bottom row: 50 μm. **b.** Ratio of fluorescence intensity of α-SMA vs. actin. (n = 4 for HPDE-WT 72 hours after seeding, n = 3 for all other treatments). Error bars indicate standard error of the mean. p-values come from one-way ANOVA tests. **c.** Bar plots of the data obtained from ELISA test IL-6 analysis for each culture condition (HFF1, HFF1+HPDE-WT and HFF1+HPDE-KRAS) grouped per time step (n ≥ 3). Each condition has been assayed in duplicate following the manufacturer’s instructions. Tukey’s multiple comparisons test: *p < 0.05, **p < 0.01, ***p < 0.001, ****p < 0.0001. **d.** Density maps of the ratio of α-SMA intensity over actin intensity in individual fibroblasts vs. distance from the cell to the edge of the duct. Data acquired from samples 7 days after printing (72 hours after seeding). Number of fibroblasts: *n_No HPD_*_E_ = 143, *n_HPDE W_* _T_ = 231, *n_HPDE KRAS_* = 350 from 2 independent experiments.

The developed model is the first to recapitulate the tumor-stroma interplay occurring in pancreatic cancer at early stages while also accurately reproducing the anatomical structure of the exocrine gland. The geometrical and morphological features of a tissue can affect the cell functionality and therefore represent another crucial aspect to consider in the design of a biomimetic model^44, 95^. Although different engineering strategies have been adopted to obtain tubular lumen structures^44–46, 83, 96^, they fail in creating the 3D tubuloacinar gland geometry exhuastively^45, 83^ or in incorporating the stromal component^46, 97^. Additional work should be performed, however, to further enhance the biomimicry of this model like incorporating other cells involved in pathology development. For instance, tissue-resident immune cells could be included inside the construct to assess the role of immune system in the early stages of pancreatic cancer progression.

In the end, this model recapitulates the tumor-associated fibroblasts activation and could consequently open new avenues to understand the role of the tumor microenvironment in pancreatic cancer progression and offer a new and relevant platform to establish effective therapeutical strategies. Our approach permits to overcome the limitations of the existing *in vitro* models that do not properly mimic the tubuloacinar geometry, the cell composition and the cell-stroma interplay of the exocrine pancreas environment. In addition, it represents a valid alternative to the costly and low-throughput animal models which are ethically questionable and limited in emulating the stromal components of PDAC^98, 99^. Indeed, the rapid fabrication process allows to obtain several scalable human models that can be tested and validated according to a high throughput screening approach.

## 5. Conclusions

In this study, we have developed a 3D *in vitro* model which mimics the complex three-dimensional microanatomy of the exocrine pancreas to study the mechanisms that take place during the early stages of pancreatic cancer. We used volumetric bioprinting to fabricate tubuloacinar gland structures in a rapid and one-step process. We showed that this biofabrication approach allows the series production of several human models with shape-fidelity, high resolution and geometrical accuracy. The GelMA-based environment proved optimal in promoting the proliferation of stromal cells which remain viable and active for several weeks within the gel structure thus permitting a long follow-up. Moreover, the construct can be monitored over time in an accessible and non-destructive way by microscopy to quantitively interrogate the model and easily obtain information.

The co-culture of human pancreatic ductal epithelial cells, overexpressing the KRAS oncogene, and stromal cells in this biofabricated *in vitro* model can recapitulate the pancreatic TME as confirmed by the stromal cells’ activation through the tumor-stroma crosstalk. In particular, we demonstrated the ability of this model in reproducing the stromal cells activation, involved in pancreatic cancer evolution, in a very short period (3 days under co-culture and 7 days after biofabrication). These results validate a scalable approach,potentially applicable in a personalized medicine workflow, in which the patients’ own cells are used to build many models of the exocrine pancreas’ microanatomy to swiftly adjust the therapy to the patient. This could enhance treatment outcomes and reduce healthcare costs. Thus, the demonstration of this fully human 3D model represents a powerful tool for the understanding of mechanisms implicated in pancreatic cancer insurgence and for developing new diagnostic and therapeutical approaches.

## 6. Methods

### GelMA Hydrogels

GelMA was produced from porcine gelatin (Sigma) following the protocol by Van De Bulcke et al.^100^. Briefly, type A porcine gelatin powder (Sigma, G2500) was fully dissolved at 10% w/v into Phosphate Buffered Saline (PBS) 1x at 50 °C. Methacrylic anhydride (Sigma, 760-93-0) was added dropwise for gelatin modification at 50 °C for 3 h. The solution was then lyophilized and stored away from light at −20 °C until use. The 5% w/v GelMA solution was created by reconstituting lyophilized GelMA powder into sterile DMEM without phenol red (ThermoFisher, 31053028) with Lithium phenyl-2,4,6-trimethylbenzoylphosphinate (LAP, Sigma-Aldrich, 900889) at a concentration of 0.16 mg mL^−1^ and filter-sterilized at 40°C. GelMA solutions were stored away from ambient light at 4 °C for no longer than 2 weeks. Before use, the rheological properties of GelMA hydrogels were evaluated employing a stress-controlled rheometer (AntonPaar GmbH, MCR302) equipped with 25 mm parallel plate geometry. In order to evaluate the photocrosslinking kinetics, filtered GelMA solution + LAP was poured on the rheometer plate and time sweep test was performed using a visible light source at 405 nm wavelength (Prizmatix, FC-LED-405A) at constant temperature (approximately 25 °C), by applying a rotational oscillation of 1 Hz and a strain amplitude of 1% in the linear viscoelastic region (measured through strain sweep test). To fabricate the more complex 3D structure, hydrogels were prepared by following the same procedure, using higher concentrations of GelMA (7% w/v) and LAP (0.5 mg mL^−1^).

#### Cell culture of HFF1

Human foreskin fibroblasts (HFF1) cells were purchased from ATCC® and cultured in Dulbecco’s Modified Eagle’s Medium (DMEM) without phenol red, supplemented with 1% Penicillin-Streptomycin (Gibco), 2% L-glutamine (Gibco) and 15% FBS (Gibco). Cells were maintained in a humidified CO_2_ incubator at 37 °C and 5% CO_2_.

#### Tomographic bioprinting

To print the proof-of-concept structure, constituted by a single acino-ductal cavity, fibroblasts were detached, counted, and centrifuged. A small volume of cells, corresponding to a final density of 0.5 million cells mL^−1^, was resuspended into GelMA with LAP and gently agitated using a 1000 μL pipette tip. 1.5 mL of the GelMA + LAP + HFF1 mix was poured into ethanol sterilized cylindrical glass vials (diameter 12 mm) with a hermetically sealing cap. All these manipulations were carried out under sterile conditions in a biosafety cabinet.

The glass flasks were dipped into water at 2°C to gel the GelMA. They were then printed using a tomographic volumetric printer. In this printer, blue light from three 405 nm laser diodes (Ushio, HL40033G) is sent through a square-core multi-mode optical fiber (CeramOptec, WF 70×70/115/200/400N), expanded, and projected on a Digital Micromirror Device (mirror size=13.7 μm, Vialux, VIS-7001), which displays the tomographic patterns. Two plano-convex lenses with focal length f_1_ = 150 mm and f_2_ = 250 mm project the images from the DMD onto the rotating vial. The vial is set to rotate using a high-precision stage (Zaber, X-RSW60C), and is inside a cubic glass container filled with cold water, acting as a refractive-index matching bath.

The calculations to produce the required tomographic patterns were performed using the software described in a previous work^50^, and scattering corrections were applied to compensate for the diffusive effects of the cell-laden hydrogels. These calculations were performed on a GPU using PyTorch^101^. This software takes 3D models in the shape of .stl files, which we designed using AutoCAD, and calculates tomographic projections using non-negative tomographic filtered back-projections. We used sets of 1000 8-bit tomographic patterns, each displayed for an angular interval of *Δθ* = 0.36°. The cylindrical vials were set to rotate at a constant angular speed of 12°/s during printing. Prints were completed in around 2.5 minutes.

After printing, glass vials were slowly heated to 27°C for 5 minutes by dipping them into water. Under sterile conditions in a biosafety cabinet, pre-warmed PBS at 37°C was gently pipetted into the glass vials, then they were gently manually agitated to rinse away the uncross-linked GelMA. The rinsed bioprinted fibroblast-laden constructs were carefully transferred to multi-well plates filled with cell medium and kept in the humidified CO_2_ incubator at 37°C.

The same manufacturing procedure described above was adopted to fabricate the hydrogel constructs with increased geometrical complexity and higher resolution. However, HFF1 were embedded in the GelMA solution at a higher density (1 million cells mL^-1^) to improve the model biomimicry.

#### Cell viability in bioprinted constructs

##### CellTiter-Blue Viability Assay

The viability of human fibroblasts embedded in the bioprinted gel constructs was analyzed by monitoring the metabolic activity through the fluorimetric resazurin reduction method (CellTiter-Blue, Promega, G8080) at 1, 2, 3, 4 and 9 days after the tomographic bioprinting process. The test was performed according to the manufacturers’ protocols. Briefly, culture medium was carefully removed, and constructs were washed with PBS (500 μL). A solution of 16% CellTiter-Blue in complete cell culture medium was prepared and added to the constructs, followed by 5-6 h incubation at 37 °C. At the end of the incubation period, 200 μL of the medium was pipetted into different wells of a 96-well plate, and fluorescence was measured from the bottom of the plate using a plate reader (BioTek) at 530 nm excitation and 590 nm emission. Fluorescence of CellTiter-Blue solutions in contact with GelMA hydrogels without cells were subtracted to avoid overestimations. Plates were covered with an adhesive film to prevent evaporation during the measurements.

##### Live/Dead Assay

Live/Dead Assay was carried out to further evaluate the HFF1 viability over the culture period, at pre-determined time points (1, 3, 7 and 14 days). Specifically, the Live/Dead solution was prepared by adding ethidium homodimer-1 (Adipogen, CDX-E0512-M001, 2mM in DMSO) and calcein-AM (Merck, 206700-1MG, resuspended to 4mM in DMSO) to PBS in concentrations of 4 μM and 2 μM respectively. The solution, prepared afresh every time it was used, was vortex-agitated for some seconds and kept at room temperature (RT) protected from light. The cellularized constructs were rinsed once with pre-warmed PBS at 37°C and transferred to a 24-well plate with wells filled with 800 μL of Live/Dead solution. Constructs were incubated in the dark for 1h, with gentle manual agitation every 15 minutes. Samples were rinsed twice with PBS and placed in optical-grade multiwell microscope slides for imaging. Imaging was conducted immediately after staining and performed in a fluorescence confocal inverted microscope (Leica, SP8) with 5x NA 0.15 (Leica, HC PL Fluotar, WD 13.7 mm), 10x NA 0.30 (Leica, HC PL Fluotar, WD 11.0 mm), and 20x NA 0.75 (Leica, HC PL APO, WD 0.62 mm) objectives. In the microscope, calcein-AM was excited at 488 nm and its emission collected from 498 to 542 nm. Ethidium homodimer was excited at 552 nm. To avoid crosstalk with the emission spectrum of calcein-AM, the emission of Ethidium was collected from 620 nm to 650 nm.

#### Epithelization of the cavity

Human pancreatic ductal epithelial cells (HPDE) stably expressing activated KRAS (HPDE-KRAS) and wild-type HPDE (HPDE-WT) were cultured in RPMI-1640 medium (Gibco, ThermoFisher Scientific) supplemented with 1% Penicillin-Streptomycin (Gibco), 1% L-glutamine (Gibco) and 10% fetal bovine serum (FBS) (Gibco). Cells were maintained in a humidified CO_2_ incubator at 37 °C and 5% CO_2_. These cells were kindly provided by Prof. Bussolino (Candiolo Cancer Institute, FPO – IRCCS, Candiolo, Italy). HPDE-KRAS cells were obtained by transducing the HPDE cell line with the oncogenic K-RasG12V as described by Siddiqui et al.^102^.

The proof-of-concept fibroblast-laden hydrogels were seeded with HPDE cells (HPDE-KRAS or HPDE-WT) by injecting them in the single acino-ductal cavity in a volume of 10 μL. After injection, the constructs were placed in 24-well plates with enough cell medium to keep them hydrated, but not enough to cover the entry of the lumen, to prevent HPDE cells from floating into the media. Two hours later, more medium was added to the wells, this time covering the full constructs.

To epithelize the multiacinar convoluted constructs, HPDE-KRAS cells were detached from the culture flask, counted and resuspended to a 7 μL volume. Then, they were manually injected with micropipette into the cavity of the fibroblast-laden hydrogels. Specifically, before the epithelization, the rinsed bioprinted fibroblast-laden constructs were cultured for 5 days. Cells were seeded at different densities (7000 cells μL^-1^ and 35000 cells μL^-1^) to test the effect of HPDE-KRAS cell density on the epithelization progress. After injection, the constructs were placed in 48-well plates and kept for 1 h at room temperature under shaking at 60 RPM.

Co-cultures were maintained in DMEM/F-12 supplemented with 15% FBS (Gibco), 1% Penicillin-Streptomycin (Gibco) and 2% L-glutamine (Gibco) since previous tests demonstrated the efficacy of this culture medium composition in promoting the cell viability^103^. the constructs were kept in a humidified CO_2_ incubator at 37 °C and 5% CO_2_.

#### Immunofluorescence microscopy

##### Immunostaining of proof-of-concept constructs

Bioprinted proof-of-concept constructs were fixed in formaldehyde 4% v/v in PBS for 5 minutes, then rinsed with PBS twice and kept at 4°C. The fixed constructs were embedded in low-melting point agarose 4% w/v in pre-warmed PBS and sliced to a thickness of 300 μm with a vibratome (Leica Biosystems, VT1000S) filled with PBS 1x. Samples were sliced orthogonally to the axis of the duct, in order to obtain circular cross-sections. After slicing, the surrounding agarose was detached gently with a brush. Slices were put onto microscope slides with adhesive imaging spacers making wells (Merck, GBL654004-100EA), covered with PBS and with a coverslip and kept at 4°C in a dark wet chamber until they were stained for imaging. Samples were permeabilized with 0.1% Triton X-100 in PBS for 10 minutes at RT, then washed 3 times for 5 minutes with PBS + 0.1% Triton X-100 (PBST) at RT. Then, samples were blocked with 2% bovine serum albumin (BSA) in PBST for 60 minutes and rinsed once with PBS. Primary antibodies, rabbit polyclonal to alpha smooth muscle actin (Abcam, ab5694-100ug, 1:50), and mouse monoclonal fibroblasts antibody TE-7 (Novus Biologicals, NBP2-50082, 1:80) in PBST + 1% BSA were incubated for 36h at 4°C. Samples were then rinsed 3 times with PBST at RT for 5 minutes. The secondary antibodies, donkey anti-rabbit IgG + Alexa 647 (ThermoFisher, A-31573) and donkey anti-Mouse IgG + Alexa 568 (ThermoFisher, A10037), were incubated at a concentration of 1:200 in PBST + 1% BSA for 2h at RT. Samples were rinsed with PBST for 5 minutes at RT 3 times. Then, ((R)-4-Hydroxy-4-methyl-Orn(FITC)⁷)-Phalloidin (1:60, 0.16 nmol mL^−1^) was incubated in PBST + 1% BSA for 30 minutes at RT. Samples were rinsed with PBS for 5 minutes at RT 3 times. They were then stained with DAPI in PBS (1:1000) for 5 minutes at RT, washed once with PBS, and finally covered with coverslips for imaging. Samples were kept in wet chambers and protected from intense light during all the immunostaining protocol.

##### Immunostaining of multiacinar convoluted constructs

The hydrogel-based VBP constructs were fixed in a 4% v/v formaldehyde solution in PBS for 20 minutes, followed by two PBS rinses and storage at 4°C. Subsequently, they were permeabilized with 0.2% Triton X-100 in PBS for 10 minutes at RT and then washed thrice for 5 minutes each with PBS containing 0.1% Tween 20 (PBST) at RT. Following this, the samples were subjected to blocking with a 2% bovine serum albumin (BSA) solution in PBST for 60 minutes, followed by a single rinse with PBS. Next, the samples were exposed to the E-cadherin monoclonal primary antibody (HECD-1; 1:2000; 13-1700, Invitrogen) in PBST supplemented with 1% BSA for 24 hours at 4°C, followed by three PBS washes at RT for 5 minutes each. The secondary antibody, Alexa 647 (ThermoFisher, A-31573), was then applied at a concentration of 1:250 in PBST with 1% BSA for 3 hours at RT, followed by three 5-minute PBST rinses at RT. Subsequently, Atto 488 Phalloidin (1:60, 0.16 nmol mL−1; 49409, Sigma-Aldrich) was incubated in PBST with 1% BSA for 30 minutes at RT, followed by three 5-minute PBS washes at RT. Finally, the samples were stained with DAPI in PBS (1:1000) for 5 minutes at RT, washed once with PBS, and covered with coverslips for imaging. Throughout the immunostaining protocol, the samples were kept in wet chambers and shielded from intense light.

##### Confocal Microscopy

Samples were imaged with a motorized inverted confocal microscope (Leica SP8) using a 10x NA 0.30 air objective (WD = 11.0 mm, HC PL Fluotar, Leica), 5x NA 0.15 air objective (WD = 13.7, HC PL Fluotar, Leica), 25x NA 0.95 water immersion objective (WD = 2.40, HC PL Fluotar, Leica). Fluorescence excitation was performed with solid-state lasers at 405, 488, 552, and 638 nm, and its emission was collected with two twin Hybrid Detectors. An additional photomultiplier tube collected transmitted light from the excitation laser. To acquire images of the full cross sections of the bioprinted constructs, the automatic motorized stage was used to take sequential images along grids that were later stitched together. Two sequential two-channel acquisitions were performed for each sample; one collecting the fluorescence from DAPI (440-480 nm) and TE-7-bound secondary antibody (568-620 nm), and another collecting the fluorescence from Phalloidin-FITC (498-542 nm) and α-SMA-bound secondary antibody (648-720 nm). Both acquisitions used the same grid coordinates of the motorized stage and included a brightfield image acquisition. Lasers’ intensities, detector gain, and optical path were kept unchanged across image acquisitions to guarantee intensities were comparable. Microscopy images were automatically acquired using LAS X software (Leica). Images were acquired for almost 100 slices of 14 independent biological samples.

##### Light sheet microscopy

After immunostaining, samples were imaged with a light sheet microscope (Z1, Zeiss) using a 20x NA 1.0 corrected water dipping objective (WD = 1.8 mm, W Plan Apochromat, Zeiss) or a 5x NA 0.16 air objective (WD = 18.5 mm, EC Plan Neofluar, Zeiss). Samples were embedded in low-melting point agarose and imaged immersed in distilled water. Excitation was performed at 405 nm, 488 nm, and 639 nm, with fluorescence from 488 and 639 nm in two-camera dual-color acquisitions (using a dichroic mirror, cut wavelength 660 nm). PCO.Edge sCMOS cooled cameras (1920 × 1920 px) were used. Resulting images were stitched with Zeiss’s Zen software and analyzed and processed with ImageJ.

##### Image processing

Due to the large area of the acquired microscopy images (> 250 mm^2^ in some cases) and to the fact that we were imaging soft, elastic hydrogels, there was displacement between the DAPI-TE-7 and the Phalloidin-α-SMA images for some samples (note however, that because Phalloidin and α-SMA were always acquired in parallel and not sequentially, there was never displacement between these two channels).

Displacement between the DAPI-Te-7 and Phalloidin-α-SMA images was corrected with a custom-made Python code. The code compared the bright-field channels of corresponding DAPI-Te-7-BF and Phalloidin-α-SMA-BF tiles, calculated the necessary homography (n*_features_* = 5000) that needed to be applied to the DAPI-Te-7-BF image so that it matched the Phalloidin-α-SMA-BF image^104^. The code would then apply such homography and save a transformed copy of DAPI-Te-7-BF image.

Images were batch-stitched together using the Grid/Collection stitching plugin^105^ on ImageJ^106^ using the Phalloidin channel as reference. Multichannel microscopy images (DAPI, Phalloidin, Te-7, α-SMA, BF) depicting multiple slices of the same bioprinted construct were then manually cropped to fit only one slice per image.

##### Inflammation quantification from fluorescence data

Inflammation was quantified from microscopy images by measuring the ratio between the intensity of α-SMA vs. actin. This calculation was done with a custom-made Python code. The code first segments cell nuclei from the DAPI channel. Then, the code segments all regions of at least 13.4 μm^2^ with a non-zero actin or α-SMA intensity in size and which are adjacent to a cell nucleus. An average intensity is calculated for these masked regions for the actin and α-SMA channels, and a ratio is reported. Actin and α-SMA intensities were additionally normalized to excitation light intensities, to make the ratio comparable across multiple acquisitions.

##### Single-cell measurements of inflammation

The ratio of actin to α-SMA intensities was also computed for segmented single cells. Hand-made digital annotations of the outline of the inner channel of the bioprinted pancreatic constructs (based on the brightfield and fluorescence channels of the microscopy images) were used to measure the distance of the individual cells to the channel.

##### Visualization of microscopy images

Multi-channel microscopy images were visualized using ImageJ and the Look-Up Tables from Christophe Leterrier (https://github.com/cleterrier/ChrisLUTs) for the Live/Dead experiments and from the BioImaging and Optics Platform at EPFL (https://biop.epfl.ch/Fiji-Update/luts/) for the co-culture inflammation experiments. Brightness and contrast were set the same for each channel of all images that were compared (particularly actin and α-SMA). Sketches of cell-laden constructs were created with BioRender.com. Plots were produced using matplotlib.org^107^ and seaborn.pydata.org^108^.

#### ELISA Assay

The concentrations of cytokines were assessed in the cell supernatants collected at 24-, 48- and 72-hours post-seeding of HPDE cells. These supernatants were obtained from wells containing the VBP constructs seeded with HFF1 cells, HFF1 cells co-cultured with HPDE-KRAS, and HFF1 cells co-cultured with HPDE-WT cells. Quantification of IL-6 cytokines was conducted using the IL-6 Human ELISA Kit (CRS-B001-96tests, ACROBiosystems). The concentrations were determined by referencing a standard curve, where absorbance values of each standard sample were plotted against the corresponding concentrations of human IL-6 standards.

#### Statistical analysis

Data were arranged and analyzed using Pandas^109^. The graph data are presented as the mean ± standard deviation (SD) for at least three independent experiments (n ≥ 3). Significance was measured with one-way ANOVA followed by pairwise comparison with Tukey’s multiple comparisons test, using GraphPad Prism 9.3.1 for metabolic activity experiment (*p < 0.05, **p < 0.01, ***p < 0.001, ****p < 0.0001) and using SciPy’s statsmodels (https://www.statsmodels.org/stable/index.html) for inflammation quantification.

## Data availability

All data from this work and the code used to process and analyze microscopy data and perform statistical analysis are freely available from the Zenodo repository https://doi.org/10.5281/zenodo.7525930.

## Acknowledgements

We thank Dr. Lely Feletti and Prof. Aleksandra Radenovic from the Laboratory of Nanoscale Biology at EPFL for their support with cell culture and for sharing cell-culture equipment. We also thank Estée Grandidier for support with cell culture. Authors thank Dr. Jessica Sordet-Dessimoz from EPFL’s Histology Core Facility for useful discussions on immunostaining and Dr. Nicolas Chiaruttini and Dr. Thierry Laroche for support with fluorescence microscopy. We also acknowledge Ragunathan Bava Ganesh from the Laboratory of Biological Network Characterization at EPFL for his support with fluorometry. We thank Cecilia Traldi and Matteo Bortolameazzi for their support with the material characterization. The authors thank the open-source tools (and their contributors) which were used in this work, including FreeCADweb.org, Inkscape.org, Python.org, PyTorch.org, Pandas.Pydata.org, and Fiji.sc.

## Contributions

V.S. and J.M.W contributed to *Data curation*, *Formal Analysis*, *Investigation*, *Methodology* and *Writing – original draft*. A.B. contributed to *Data Curation* and *Investigation*. G.C. contributed to *Supervision* and *Writing – review & editing*. C.T.T. contributed to *Methodology*, *Supervision* and *Writing – review & editing*. C.M. contributed to *Supervision, Funding Acquisition* and *Writing – review & editing*.

