## Supplementary figures and information for "3D *in vitro* modeling of the exocrine pancreatic unit using tomographic volumetric bioprinting"

### Supplementary materials

*Properties of GelMA hydrogels*

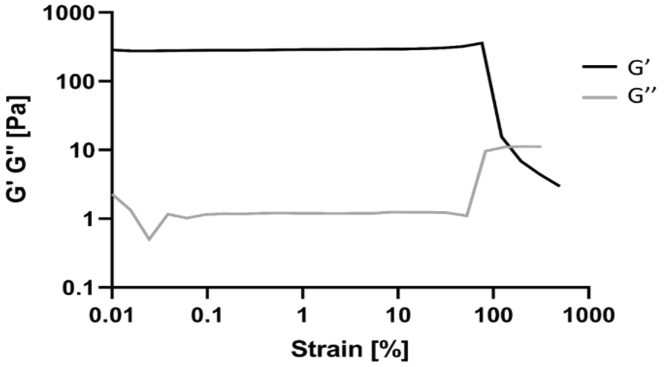

**Sup. fig. 1. Rheological properties of GelMA hydrogels.** Strain sweep test: G' (black) and G'' (gray) versus strain amplitude. Filtered solution of 5% GelMA with LAP at a concentration of 0.16 mg $mL^{-1}$ was used for the tests.

*Swelling*

| Sample number | Mass of lyophilized GelMA (g) | Mass of wet GelMA (g) | Swelling capacity (%) |
|---|---|---|---|
| 1 | 0.971 | 7.616 | 684 |
| 2 | 0.946 | 7.729 | 717 |
| 3 | 0.818 | 7.159 | 775 |
| | | **Mean Swelling capacity (%)** | 726±27 |

**Sup. table 1. Swelling capacity of GelMA hydrogels.** Swelling capacity was measured by immersing lyophilized GelMA in PBS 1x at RT for 30 minutes and then weighing the sieved hydrated GelMA with a precision scale.

Swelling capacity was measured by hydrating lyophilized weighted masses of GelMA in deionized $H_2O$ at room temperature for 24h. Then, we recovered them with an aluminum strainer and weighted them for a second time. Mean swelling capacity was calculated according to the formula:

$$\%\,Swelling\ capacity\ = \frac{Weight_{hydrated} - Weight_{dry}}{Weight_{dry}} \times 100$$

*Live/Dead Imaging*

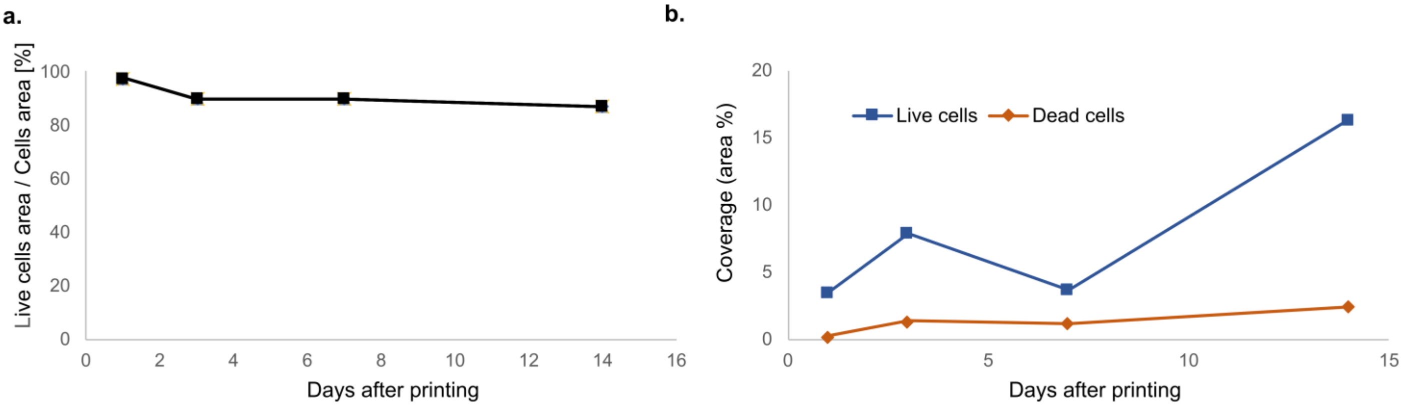

**Sup. fig. 2. Live/Dead Imaging (HFF1) in the bioprinted hydrogels. a.** Area of live cells as a percentage of the total area of cells. **b**. Area percentage coverage (area of cells/area of hydrogel) for live and dead cells. n = 1 sample per time point.

Viability was also measured from fluorescence microscopy images after a Live/Dead staining with calcein-AM and ethidium homodimer-1, as described in the methods section. The fluorescence images (area > 15 $mm^2$) were then analyzed with ImageJ. First, a maximum intensity z-projection along roughly 200 µm of gel was performed (slicing = 20 µm). Then, each of the channels was binarized. The areas of live and dead cells were then measured automatically. To measure coverage percentages, the hollow region of the duct was not computed in the hydrogel area.

These measurements suggest that fibroblasts stay viable and tend to increase their surface in the two weeks after printing.

*Cell density and epithelialization in the 3D complex VBP model.*

Different cell densities were tested to determine the optimal number of HPDE-KRAS cells required to achieve homogeneous coverage of the acini surfaces within the complex 3D construct (Fig. sup. 2). Results indicated that cells seeded at a lower density (1400 cells $mm^{-2}$; 0.9:1 HPDE:HFF1) formed cell clusters after 3 days and subsequently epithelialized most of the cavity within 3 weeks. In contrast, the same tendency was not observed with HPDE-KRAS cells seeded at a higher density (7200 cells $mm^{-2}$; 4.6:1 HPDE:HFF1). Indeed, a higher number of cells resulted in the development of cell sedimentation, distinguishable by dark spots in brightfield microscope images, at the bottom of the acini within 3 days of seeding. However, epithelial cells were observed to migrate from the sedimentation and colonize the lateral and more vertical parts of the cavity, ultimately forming a thin epithelium after 7 days.

a.

| cells mm$^{-2}$ | cell clusters | sedimentation | time until coverage > 60% |
|---|---|---|---|
| 1400* | yes | no | 21 days |
| 7200* | no | yes | 7 days |

(*) Seeding cell density

b.

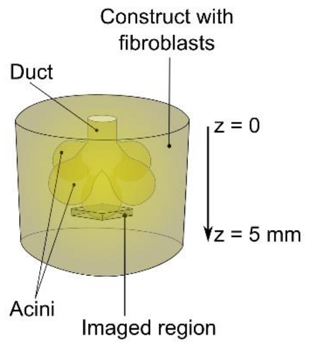

c.

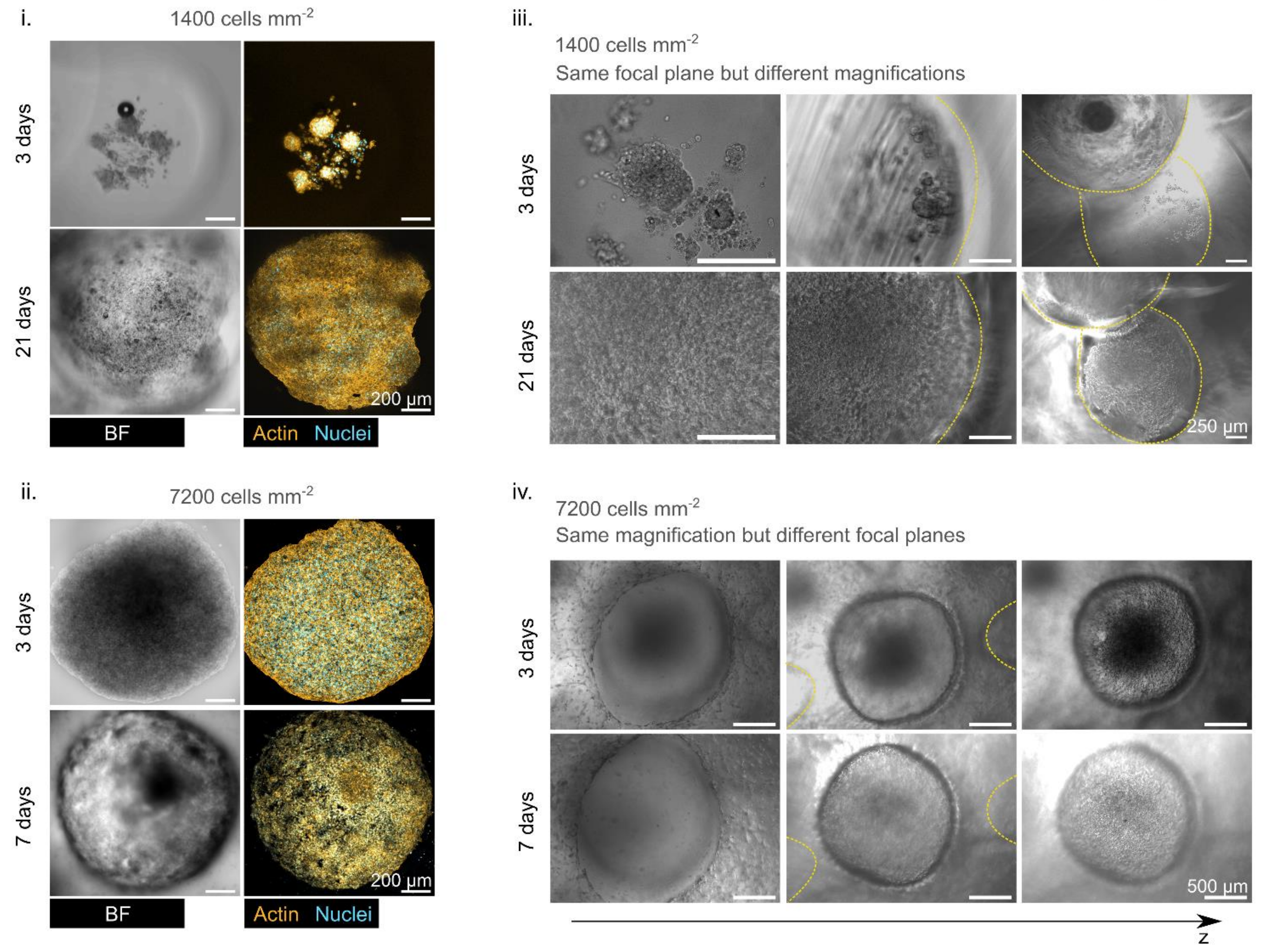

**Sup. fig. 3. Cell density in epithelization of 3D complex VBP model**. **a.** Table summarizing the behavior of epithelial cells at different cell densities. **b.** Schematic representation of the region where microscopy images were acquired. **c.** Confocal microscopy images depicting the bottom acinus at 3 and 7 days (i) and 3 and 21 days (ii) post-seeding of HPDE-KRAS cells in the VBP construct at densities of 1400 cells mm$^{-2}$ (i) and 7200 cells mm$^{-2}$ (ii). Cells seeded at lower density formed clusters after 3 days and subsequently epithelialized most of the cavity (>60%) within 21 days (iii). Conversely, cells seeded at higher density appeared sedimented at the bottom of the acini after 3 days, but they colonized lateral and more vertical parts of the cavity, ultimately forming a thin epithelium after 7 days (iv).

*Surface area and volume of the construct*

| *Surface area of duct* | *Volume of duct* | *Volume of bulk* |
|---|---|---|
| *36.8 mm$^2$* | *11.4 mm$^3$* | *53.7 mm$^3$* |

*Epithelium development on the acini inner surface*

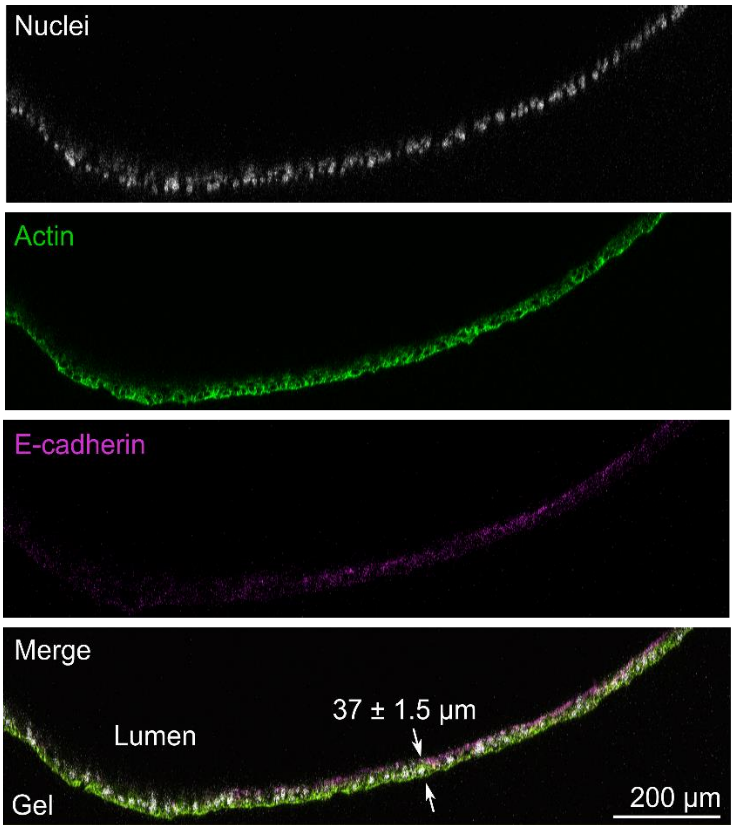

**Sup. fig. 4. Interaction between stromal and epithelial cells in VBP model**. Immunofluorescence micrographs showing the spatial distribution of E-cadherin vs. actin expression in ductal epithelial cells, from high-resolution confocal fluorescence microscopy images. The orthogonal view of this set of images was used to measure the average thickness of the resulting epithelium at 37 ± 1.5 µm. (n = 5 measurements, with standard error of the mean). Gravity pulls towards the bottom of the orthogonal view. From HPDE-KRAS in gels without fibroblasts, 14 days after seeding.

*Interaction between stromal and epithelial cells in VBP model*

When co-cultured with fibroblasts, HPDE cells still form large linings and invade the bulk in grape-like clusters as shown with arrows in sup. fig. 3a, 3b. In sup. fig. 3c multi-channel fluorescence microscopy images of the duct wall of a fibroblast-laden construct (5 days after printing process) show HFF1 assuming their characteristic elongated shape within the bioprinted structure revealing the colonization of the whole constructs by viable cells. Moreover, fibroblasts closer to the duct (where the HPDE-KRAS cells laid) qualitatively show higher α-SMA expression.

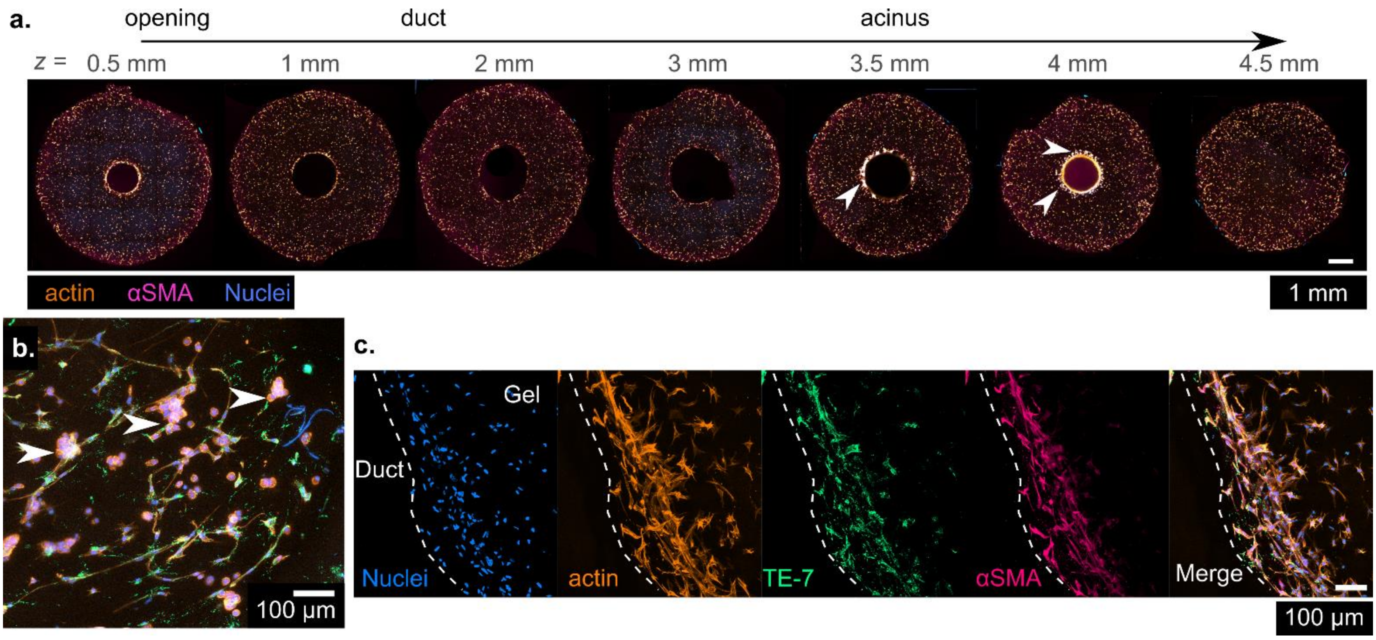

**Sup. fig. 5. Interaction between stromal and epithelial cells in VBP model**. **a.** Injected HPDE cells progress over time to line the inner face of the 3D bioprinted pancreatic model duct (Live/Dead assay). **b.** HPDE cells, shown with arrows, form grape-like clusters of smaller round cells when co-cultured with HFF1. **c.** Immunofluorescence micrographs of seven 300 μm thick slices of a 3D bioprinted pancreatic model taken 72 hours after HPDE-KRAS cells seeding and 7 days after bioprinting process. White arrows highlight migration of HPDE-KRAS cells inside the fibroblast-laden hydrogel.

**Sup. video 1. Injection of a blue glycerol-based dye into the acino-ductal cavity of a bioprinted pancreatic duct model (proof-of-concept structure).**

**Sup. video 2. VBP model (multiacinar convoluted structure).**

**Sup. video 3. 3D rotational projections from confocal microscopy images of the fibroblast-laden construct, 12 days after volumetric bioprinting.**

**Sup. video 4. 3D rotational projections from confocal microscopy images of the full epithelized construct, 7 days after HPDE-KRAS cells seeding.**
